# Flexible Learning and Re-ordering of Context-dependent Object Sequences in Nonhuman Primates

**DOI:** 10.1101/2024.11.24.625056

**Authors:** Xuan Wen, Adam Neumann, Seema Dhungana, Thilo Womelsdorf

## Abstract

Intelligent behavior involves mentally arranging learned information in novel ways and is particularly well developed in humans. While nonhuman primates (NHP) will learn to arrange new items in complex serial order and re-arrange neighboring items within that order, it has remained contentious whether they are capable to re-assign items more flexibly to non-adjacent positions. Such mental re-indexing is facilitated by inferring the latent temporal structure of experiences as opposed to learning serial chains of item-item associations. Here, we tested the ability for flexible mental re-indexing in rhesus macaques. Subjects learned to serially order five objects. A change of the background context indicated when the object order changed, probing the subjects to mentally re-arrange objects to non-adjacent positions of the learned serial structure. Subjects successfully used the context cue to pro-actively re-index items to new, non-adjacent positions. Mental re-indexing was more likely when the initial order had been learned at a higher level, improved with more experience of the re-indexing rule and correlated with working memory performance in a delayed match-to-sample task. These findings suggest that NHPs inferred the latent serial structure of experiences beyond a chaining of item-item associations and mentally re-arrange items within that structure. The pattern of results indicates that NHPs form non-spatial cognitive maps of their experiences, which is a hallmark for flexible mental operations in many serially ordered behaviors including communication, counting or foraging.

## Introduction

Mental flexibility refers to the ability to arrange thoughts or actions in novel ways. Flexible mental operations are required for many serially ordered behaviors including communication, counting, problem solving, and foraging (Davis and Pérusse, 1988; McNamee et al., 2021). A commonality of these serial behaviors is that they become more complex and flexible when subjects are able to infer the latent temporal structure on top of which mental operations can be performed. The ability to infer latent temporal structure and to mentally re-arrange representations within that structure underlies measures of intelligence and is particularly well developed in humans compared to nonhuman primates (NHP) (Dehaene, 2021; Passingham, 2021). It has remained debated, however, how capable NHPs are to infer abstract temporal structure and whether they use it to flexibly guide mental operations (Dehaene et al., 2015; Whittington et al., 2022; Passingham and Lau, 2023).

Powerful paradigms for testing how subjects infer latent structure from their experiences involves the serial ordering or items (Conway and Christiansen, 2001; Terrace, 2005). Studies using serial-order learning paradigms have shown that NHPs understand the relative ordinal position of objects in multi-item sequences (Damato and Colombo, 1989, 1990; Chen et al., 1997; Orlov et al., 2000; Jensen et al., 2019; Mione et al., 2020; Ferhat et al., 2022; Jensen et al., 2022), are able to swap object positions in a sequence when they are adjacent to each other (Matsuzawa, 1985; Scarf et al., 2011), and reverse play 3-item sequences (Xie et al., 2022; Tian et al., 2024). While these abilities highlight that NHPs show mental flexibility of manipulating rank ordered items, they also point to limitations of NHPs, when compared to humans, to extract the latent temporal structure from item sequences beyond serial chaining and rank ordering item-item associations (Dehaene et al., 2015; Zhang et al., 2022). For example, the ability to reverse play a sequence A-B-C as C-B-A involves the re-indexing of items to a new positions in a sequences (Tian et al., 2024), but this re-indexing can be achieved by swapping the relative rank of adjacent items in a sequence without the need to represent a abstract temporal structure or ordinal positions to which items are flexibly assigned (Kao et al., 2020).

Here, we set out to test the ability of NHPs for mentally re-indexing items that are non-adjacent to each other in 5-object sequences. NHPs learned sequences of objects A-B-C-D-E and were probed to re-order non-adjacent items B and D to a novel A-D-C-B-E sequence. The re-ordering of objects to different temporal positions can be achieved by independently representing the specific object items and the latent ordinal structure to which the items could be assigned (Tian et al., 2024). In computational models the latent temporal structure of experienced environments can be inferred and represented as a non-spatial cognitive map of item locations that enables the flexible re-indexing of objects to different positions on this cognitive map (Behrens et al., 2018; Whittington et al., 2022).

We tested the learning and flexible re-ordering of object sequences in four rhesus macaques using 3-D rendered objects shown on a touchscreen Kiosk station in their home cage (Womelsdorf et al., 2021). The paradigm required NHPs to choose simultaneously presented objects in a predetermined sequential order in trials that allowed maximally fifteen choices to complete the sequence and receive fluid reward (**Fig. 1A**). When a sequence was completed, or the maximum of fifteen choices was reached the subjects were presented with a new trial in which the same objects were arranged at new locations to prevent a spatial strategy. A sequence was shown for a total of fifteen trials. Subjects received for each correct object choice immediate visual feedback (yellow halo; for incorrect choice: grey halo) and an increment (for incorrect choices: a reset) of the slider position that signaled how many steps away subjects were from receiving fluid reward (see Fig. 1A and **Suppl. Video**). After fifteen trials with the same sequence, we changed the background (context 1) to a new background (context 2) that showed the same objects but required a new pre-determined sequential order with objects B and D swapping position. Each pair of context 1 / context 2 sequences used a unique set of objects. Each session evaluated multiple sequence pairs in an early block, followed by a delayed match-to-sample task, and followed by a late block of multiple sequence pairs (**Fig. 1B**).

**Fig. 1.**
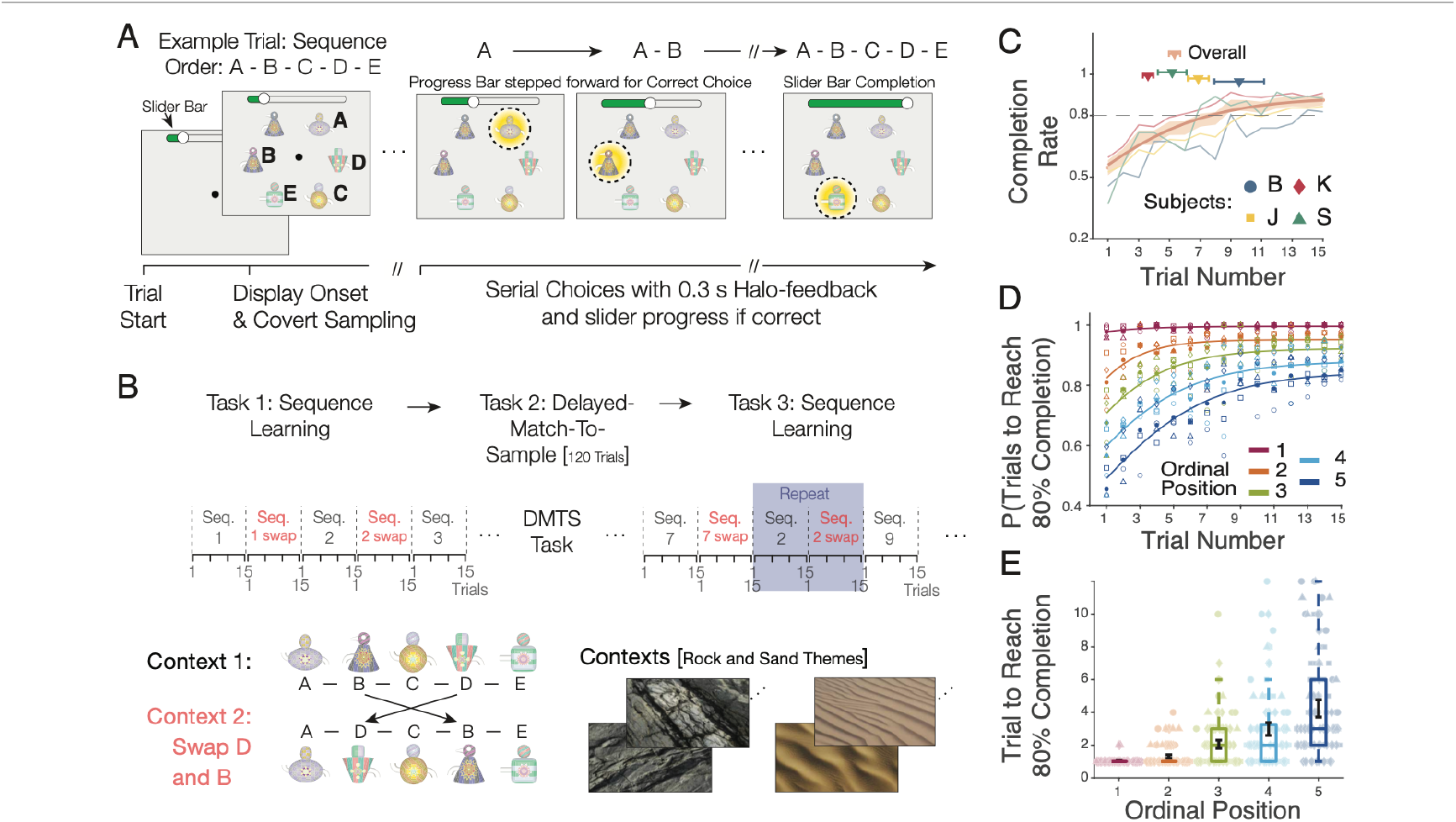
Learning 5-object sequences. (**A**) Each trial presented six objects. Monkeys learned to touch five objects in a pre-determined order A-B-C-D-E and avoid a distractor object. A correct choice led to visual feedback (yellow halo) and incremented a slider progress bar on top of the screen. Monkeys received fluid reward upon completion of the sequence. (**B**) Each sequence was presented in 15 separate trials (each with random spatial arrangement) and was followed by a sequence with the same objects but a new context background and swapped order of object B and D (*red* font). After 5-6 pairs of sequences subjects performed a delayed match-to-sample (DMTS) task for 120 trials. Then, sequence learning was performed again with ∼3 new and ∼3 repeat-sequence pairs from earlier in the session. Sequences were presented on rock- or sand-themed contexts (*see* methods). (**C**) Learning curves for completing sequences with < 10 errors (y-axis). Symbols mark the average trial (±SE) at which 80% completion was achieved consistently across all subsequent trials. (**D,E**) Probability to reach 80% correct choices (*y-axis*) for each ordinal position (diff. colors) across trials (*D*) and on average (*E*). Error bars are SE.

We found that subjects learned quickly to complete 5-object sequences A-B-C-D-E. When the background context changed to indicate that the same objects are re-ordered to A-D-C-B-E they on average anticipated the swapped positions of objects B and D and chose object D at the second ordinal position (**Fig. 1C**). This pro-active swapping was more likely when the initial sequence was learned better and correlated with working memory performance on a session-by-session level. The findings suggest that NHPs effectively use context cues to mentally re-index objects to a latent temporal structure.

## Results

### Fast learning of object sequences

We tested sequence learning in four NHP in 31.25 experimental sessions (subjects B: 13; J: 40; K: 59; S: 13). Subjects learned to perform the 5-object sequences with ten or less erroneous choices within 5.34 ± 0.37 trials (Subject B: 9.55 ± 1.62; J: 6.88 ± 0.68; K: 3.61 ± 0.34; S: 5.18 ± 0.93) (**Fig. 1C**). Across sequences, subjects successfully learned 88% of sequences above the completion criterion of 80% (Subject B: 80; J: 87; K: 93; S: 92) (**Suppl. Fig. S1A,B**), reaching above-chance accuracy at each ordinal location (**Suppl. Fig. S1C,D**). Learning gradually progressed through the ordinal positions, reaching 80% completion of the first ordinal position on average after 1.03 trials (± 0.03, 95% confidence interval), and of the 2^nd^-5^th^ ordinal position on average after trials 1.30 (± 0.11), 2.06 (± 0.25), 2.98 (± 0.39), and 4.24 (± 0.53) (**Fig. 1D, E**). Learning was achieved by reducing erroneous choices of objects, while errors indicating perseverative tendencies or violating the task rules were infrequent throughout (**Suppl. Fig. S1E,F**). Reaction times gradually increased with ordinal positions, consistent with prior studies (**Suppl. Fig. S2**) (Colombo et al., 1993).

### Subjects pro-actively re-order non-adjacent objects of well-learned object sequences

We next tested whether subjects could re-assign objects of the learned A-B-C-D-E sequence to different positions. When subjects completed 15 trials of the initial sequence, we changed the context background to a new context 2 and re-ordered the same objects to the new order A-D-C-B-E. The sequence in context 2 swapped positions of objects B and D (**Fig. 2A**, see **Suppl. Videos**). Subjects successfully used the context change as a cue and adjusted to the swapping in context 2, reaching 80% completion rate on average at trial 2.91 (± 0.61, Mean ± 95% CI), compared to 5.34 (± 0.73) trials for learning the initial sequence in context 1 (**Fig. 2B**). The faster learning of sequences in context 2 was evident in each subject (completion rate for sequences of context 1 / 2 in subject B: 5.38 (± 0.83) / 3.85 (± 0.73); J: 4.71 (± 0.53) / 2.67 (± 0.31); K: 2.98 (± 0.26) / 1.97 (± 0.17); S 5.38 (± 0.87) / 3.08 (± 0.52) (**Fig. 2C**). Reaction times were similar across ordinal positions for context 1 and 2 (**Suppl. Fig. S2**).

**Fig. 2.**
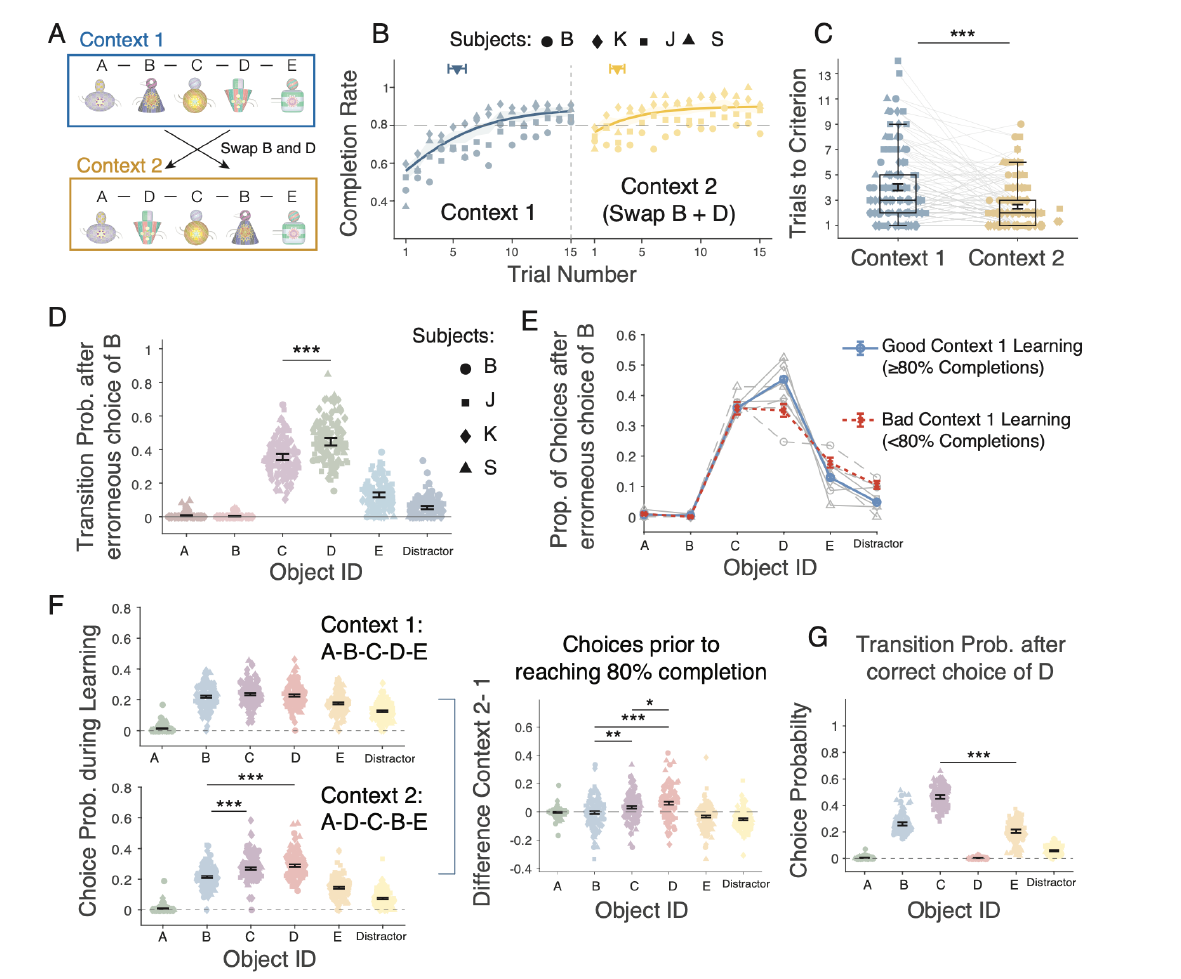
Subjects swap objects between non-adjacent ordinal positions. (**A**) Context 2 swapped the ordinal position of object B and D. (**B**) Subjects reached 80% completion rate earlier in context 2 (yellow). (**C**) Learning speed (trials to reach 80% completion) in context 1 (initial sequence) and context 2 (swapped). (**D**) Retro-active swapping after error on B: Probability of choosing objects immediately after erroneously choosing object B in context 2: for A: 0.008 ± 0.003l; B: 0.004 ± 0.002 (95% CI); C: 0.356 ± 0.019; D: 0.447 ± 0.023; E: 0.131 ± 0.015; Distractor: 0.055 ± 0.010. Three stars denote p < 0.001 difference (Welch’s t-test). (**E**) Retro-active swapping by choosing object D after an error on B was more likely when the context 1 sequence was learned at high >80% completion rate. (**F**) Choice probability for objects in context 1 (*upper*), context 2 (*lower*) and their differences (*right* panel) in trials prior to reaching the 80% completion rate. Subject more likely chose D over C and B, suggesting they pro-actively swapped object D into the 2^nd^ ordinal position. (**G**) Choice probability of objects after correctly choosing object D in context 2, showing that subjects correctly continued with C rather than E. Stars denote significance levels (Welch’s t-tests). Error bars are SEs.

How did the subjects adjust to the swapped positions of object B and D in context 2? A serial chaining framework predicts subjects will erroneously choose object B at the second ordinal position and adjust to this error by choosing the next neighboring object C at the third ordinal position because its relative position is closest to the second position. In contrast to such a serial inference, subjects may also at the second ordinal position pro-actively choose object D. This would reveal they understood that D was indexed to the absolute fourth position in context 1 and needed to be re-arranged to the new A-D-C-B-E sequence when the context changed. We found that overall, subjects re-indexed object D from the fourth ordinal position in context 1 to the second ordinal position in context 2 more likely than choosing object C, which occupied the adjacent thurd position in context 1. First, we analyzed choices in context 2 prior to reaching completion rate. When subjects erroneously choose object B in context 2 they were more likely to correctly choose object D rather than object C (Welch’s t-test, p = 6.2 × 10^-9) (**Fig. 2D**). This correct re-indexing of object D in context 2 was more likely when the initial A-B-C-D-E sequence in context 1 was learned at ≥ 80% completion rate (**Fig. 2E**). When context 1 was performed below 80% completion rate, subjects similarly often applied the correct re-indexing strategy (choosing object D at the second position) and the incorrect serial-inference strategy (choosing object C at the second position) (**Fig. 2E**).

We next quantified directly whether subjects used the context cue to swap objects pro-actively, i.e. whether - after correctly choosing A - they anticipated choosing object D at the second ordinal position in context 2 without erroneously choosing object B. We found that compared to context 1, in context 2 subjects more likely choose object D as well as object C more likely than B prior to reaching an 80% completion rate (t-test, p < 0.001; **Fig. 2F**). Directly comparing contect 1 versus context 2 showed that already prior to reaching 80% completion choosing D (indicating swapping) was significantly more likely than choosing C (indicating serial inference) (t-test, p = 0.05; **Fig. 2F**). This pattern of results suggests that subjects considered both, object C and object D as possible target objects in context 2 early during learning when they had not yet achieved an 80% completion rate, but that they more likely anticipated that object D was the most likely object to be correct at the second ordinal position.

To further test whether subjects considered object D to be the correct object swapped into the second ordinal position in context 2 we calculated how likely they chose object C (correctly) rather than object E (incorrectly) following correctly choosing D. Subjects chose C more likely than object E following correctly choosing D in the second context, indicating they swapped D to an earlier position rather than jumping to the late chain of D-E sequence they learned in context 1 (**Fig. 2H**) (Welch’s t-test, p < 1 × 10^-10; correctly choosing object C: 0.465 ± 0.014, incorrectly choosing object E: 0.206 ± 0.014 and object B: 0.261 ± 0.013).

### Sequence memory and rule memory improves swapping ability

To swap the position of objects in a sequence depends on recalling the original sequence from memory and on an understanding of the swapping rule. To test how long-term memory affected swapping performance we split each experimental session into early sequence learning, followed by an interspersed delayed-match-to-sample task, and late sequence learning (**Fig. 1B**). During the late learning epoch, we repeated a subset of sequence pairs from the early epoch and tested whether the early sequence was memorized and potentially facilitated performance and the likelihood of swapping of the repeated sequences in the late epoch. We found that repeated sequences were learned faster than the initial sequence shown earlier in the session, or other new sequences that were interspersed late in the session to control for a possible effect of the time-in-task (trials to 80% completion: Early New sequences: 4.12 (± 0.63; Mean ± 95% CI); Late New sequences 4.74 (± 0.80); Repeated sequences: 1.81 (± 0.37), **Fig. 3A**; **Suppl. Fig. 3**). The likelihood of pro-active swapping was already above chance for the early sequences and did not increase further in repeated sequences (New Early vs. Repeat: p = 0.0818; New Early vs. New Late: p = 0.0998; Repeat vs. New Late: p = 0.6962; **Fig. 3B**). However, in repeated sequences subjects were significantly more likely in context 2 to retro-actively swap object D into the second position after erroneously choosing object B at the second ordinal position (New Early vs. Repeat: p = 0.0008; New Early vs. New Late: p = 0.0019; Repeat vs. New Late: p = 0.3462; **Fig. 3C**). Thus, sequence memory improved error correction in trials when subjects failed to pro-actively swap object D into the second ordinal position.

**Fig. 3.**
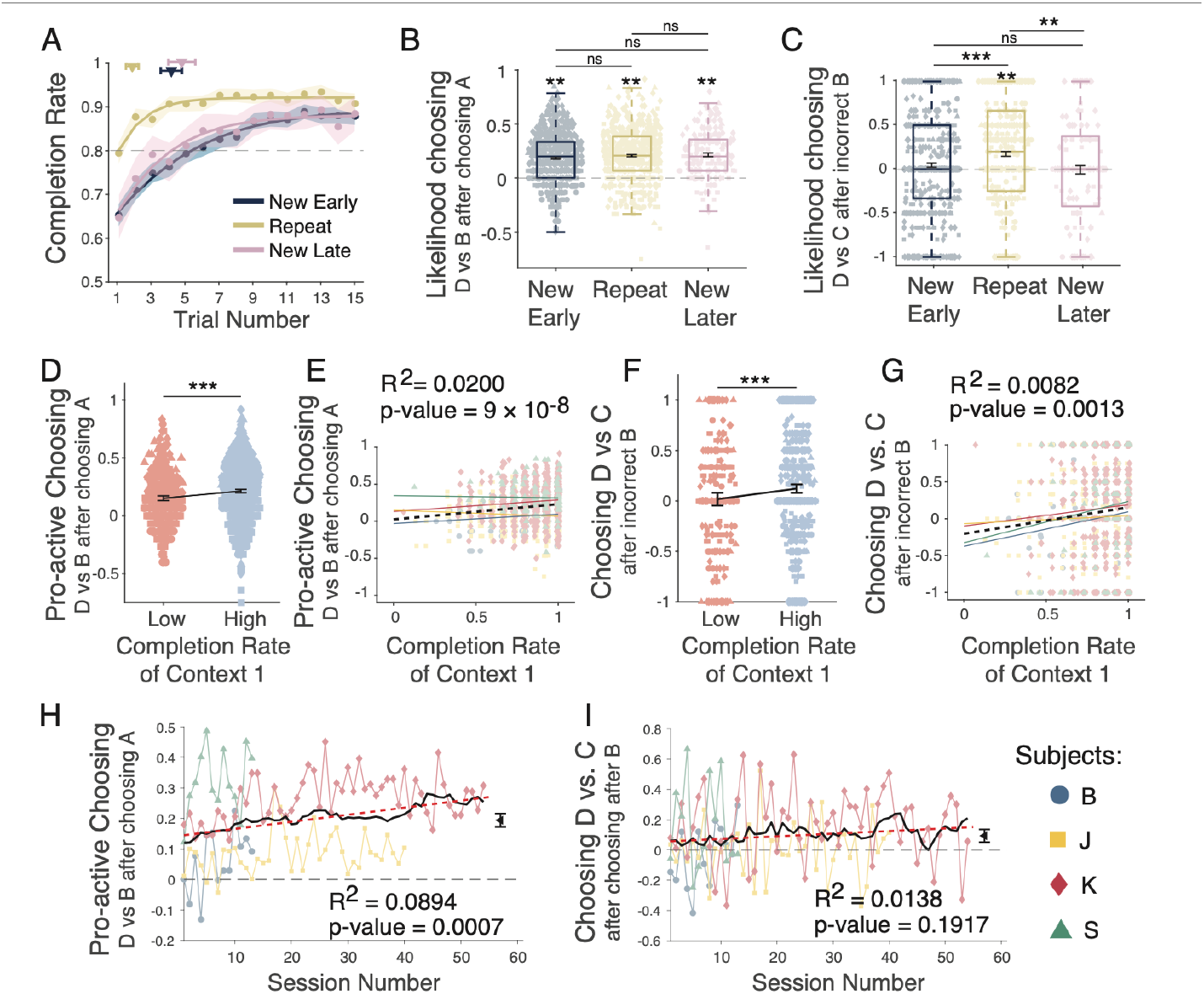
Memory of object sequences improves pro-active and retro-active swapping performance. (**A**) Completion rate of sequences before the working memory task (New Early), and for new and repeated sequences performed after the working memory task (New Late, Repeat). (**B**) Pro-active swapping in context 2: Anticipating that D is in 2^nd^ position in context 2 does not differ between conditions. (**C**) Retro-active swapping after erroneously choosing object B in context 2: Difference of choosing object D vs. C. (**D**) Better initial performance in context 1 (blue *vs* red: higher *vs* lower completion rate) is associated with higher likelihood of pro-actively swapping object D and B context 2. (Low completion rate: M = 0.1525, 95% CI [0.1315, 0.1736]; high completion rate: M: 0.2148, 95% CI [0.1999, 0.2297]; p: 2.58 × 10^-6). (**E**) Regression of the completion rate in context 1 and the likelihood to choose object D immediately after object A in context 2 (β: 0.3568; intercept: - 0.2040; R^2^: 0.0082; p: 0.0013; Cohen’s f^2^: 0.0082). (**F**) Same format as *D* for retro-active swapping, i.e. for choosing object D in context 2 at the 2^nd^ ordinal position after erroneously choosing B (low completion rate: M: 0.0179, 95% CI [-0.0458, 0.0815]; high completion rate: M: 0.1226, 95% CI [0.0809, 0.1643]; t-test, p: 0.00714). (**G**) Regression of context 1 completion rate and retro-active swapping in context 2 (β: 0.2073; intercept: 0.0249; R^2^: 0.02; p: 8.98 × 10^-8; Cohen’s f^2^: 0.0204). (**H**) Choosing object D proactively after object A in context 2 was more likely than choosing object B on average across 124 valid sessions (M: 0.19, 95% CI [0.17, 0.21]; p < 1 × 10^-10). The effect increased in strength over sessions (regression β value: 0.0023; p: 0.0007; Cohen’s f^2^: 0.0982). (**I**) Choosing object D after an erroneous choice of B in context 2 was more likely than choosing object C on average across 124 sessions (M: 0.09; 95% CI [0.05, 0.13], one-sample t-test, p = 2.5 × 10^-5). The regression slope across sessions (red dashed line) was not significant (β value: 0.0017; p: 0.1917; Cohen’s f^2^: 0.0140).

Next, we analyzed whether the depth of memorizing the initial sequence early in the session influenced the swapping performance of that sequence later in the session. We found that subjects more likely pro-actively swapped D into the second position in context 2 for repeated sequences late in the session when the sequences were better memorized early in the session, i.e. when they were performed at a higher completion rate (R^2^: 0.02, p=9×10^−8^) (**Fig. 3 D,E**). Similarly, retro-active swapping of object D after erroneously choosing B in context 2 at the second position was also more likely late in the session when the initial sequence was better learned early in the session (R^2^: 0.0082, p=0.0013) (**Fig. 3 F,G**).

In addition to memory for the sequence, subjects may also improve memory of the swapping-rule that required swapping the 2^nd^ and 4^th^ ordinal position once the context background changed. We analyzed rule memory by quantifying how pro-active and retro-active swapping changed over sessions. Pro-active swapping was apparent already in the first experimental sessions and gradually increased over sessions (R^2^: 0.0089, p=0.0007; **Fig. 3F**), while the likelihood of retro-active swapping remained similar across sessions (R^2^: 0.0138, p=0.1917) (**Fig. 3I**).

### Pro-active swapping of object positions correlates with working memory

Swapping objects from later and earlier ordinal positions may involve transiently storing the position index of the original objects in a temporary variable, which is similar to a short-term memory buffer (Tian et al., 2024). We thus hypothesized that working memory ability predicts pro-active swapping performance, which we tested by assessing delayed match-to-sample performance in the same behavioral sessions that assessed swapping (**Fig. 4A, Suppl. Fig. S4**). We found that WM performance did not correlate with overall sequence learning accuracy (**Fig. 4B**), but WM accuracy significantly correlated with pro-active swapping abilities on a session-by-session basis (**Fig. 4C**). This result indicates that successfully swapping object D and B in context 2 is not only influenced by longer term sequence-memory and rule-memory (**Fig. 3**), but also by an ability to hold objects active in short-term working memory.

**Fig. 4.**
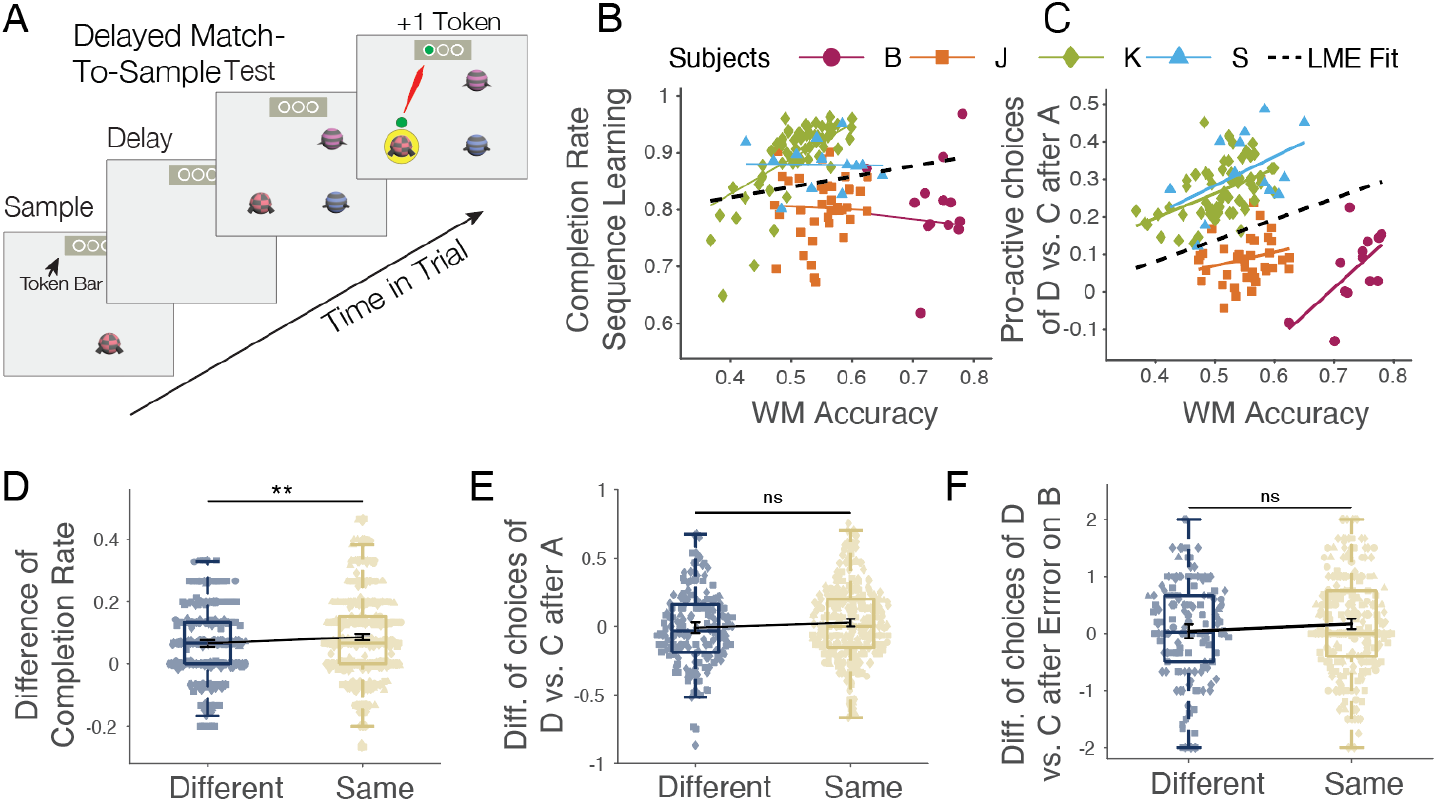
Working memory performance correlates with pro-active swapping. (**A**) Delayed match-to-sample (DMTS) paradigm. (**B**) Across sessions, the DMTS accuracy did not correlate with the completion likelihood of sequences. (**C**) DMTS accuracy significantly correlated with pro-active swapping, i.e. with choosing object D after A in context 2. Black line denotes avg. Linear Mixed Effect model; colors show individual subjects. (**D**) Contexts facilitated sequence learning: Completion rates were higher for sequences repeated with the same context (n=687, *yellow*) than a different context (n=430, *blue*): (same contexts: 0.0865 (95% CI [0.0763, 0.0967]; different contexts: 0.0660, 95% CI [0.0549, 0.0771]). (**E**) Context did not modulate pro-active swapping (different context: M = -0.0093, 95% CI [-0.0363, 0.0177]; same context: M = 0.0373, 95% CI [0.0095, 0.0651]; Welch’s t-test: p = 0.1269). (**F**) Context did not modulate retro-active swapping (different context: M = 0.0386, 95% CI [-0.1273, 0.2045]; same context: M = 0.1225, 95% CI [0.0319, 0.2131]; Welch’s t-test: p = 0.3001).

We next tested whether memory of the context background in which a sequence was learned influenced swapping behavior. When a sequence was repeated after the working memory task, it was presented on either the same, or on a different contextual background than the initial new sequence. We found improved sequence learning performance on repeated versus early (initial) sequences when the context was the same than different (Welch’s t-test, p = 0.0079; **Fig. 4D**). However, this overall contextual facilitation did not modulate pro-active or retro-active swapping in context 2 (pro-active sapping: Welch’s t-test: p = 0.1269; **Fig. 4E**; retro-active swapping: Welch’s t-test: p = 0.3001; **Fig. 4F**).

## Discussion

We found that rhesus monkeys learned multiple 5-object sequences in single experimental sessions (**Fig. 1, Suppl. Video**). When a change in context indicated that non-adjacent objects B and D switched positions, subjects swapped these objects more likely than choosing the next-ranked object C in the sequence (**Fig. 2D-G**). The swapping occurred pro-actively, i.e. prior to making an error (**Fig. 3B,D,H**), as well as retro-actively, i.e. when subjects corrected an erroneous choice of B at the second ordinal position in context 2 (**Fig. 3C,E,I**). Four factors were associated with better swapping performance: Swapping in context 2 was more likely when the sequence had been learned at a higher proficiency level in the immediately preceding context 1 (**Fig. 2E**) and in repeated sequences in context 2 when context 1 was performed at higher proficiency level ∼30-60 min prior in the early epoch of a session (**Fig. 3D-G**); when the swapping rule had been performed for more sessions later in the experiment (**Fig. H**); and when subjects showed a higher working memory performance (**Fig. 4C**). Taken together, these results show that rhesus monkeys infer the latent temporal order of objects during sequence learning and are able to flexibly swap the index of object identities to absolute positions in that latent order when a new context instructs them to reassign objects to new temporal positions.

### Swapping behavior reflects flexible mental re-indexing of object associations

Successful pro-active swapping behavior shows that monkeys used the context of the swapped sequence as a cue to re-order objects A-B-C-D-E of context 1 to a new A-D-C-B-D order in context 2. A neuronal correlate for such cue-triggered reconfiguration process has recently been suggested in an experiment that required NHPs to reverse play the spatial order of 3-item sequences (Tian et al., 2024). Groups of frontal cortex neurons represented forward spatial 3-item sequences A-B-C and re-coded the sequence when a visual cue required to report the sequence backwards. During the rec-coding process neuronal population activity in frontal cortex transiently encoded the swapped item positions, which was followed by a new neuronal population that encoded the backward sequence (Tian et al., 2024). While these neuronal findings were limited to reversing adjacent items, we suggest that they provide a versatile framework to conceptualize the re-indexing of objects to non-adjacent positions in our study. In particular, this framework predicts that in our study objects of the A-B-C-D-E sequence in context 1 will be encoded by neurons indexing their ordinal position (Xie et al., 2022). When the context changes and objects D and B need to be swapped, the original index of objects B and D are temporarily encoded in a short-term buffer, and the original A-B-C-D-E sequence is reconfigured into the new A-D-C-B-E sequence. When this operation completes, the transient buffer is not needed anymore and the swapped sequence is encoded by a group of neurons that is partially distinct from the group of neurons that encoded the initial sequence (Tian et al., 2024). The key insight of this framework is that the swapping operation can be conceived of as the re-indexing of objects to the absolute serial position of a sequence. This indexing operation requires that the objects and the temporal sequential structure are independently encoded in the neuronal network that performs this operation, which is well supported by neurophysiological evidence (Xie et al., 2022; Tian et al., 2024). Our study suggests that NHPs uses these neuronal processes when learning to re-index objects to non-adjacent positions, which extends previous findings and documents a high level of flexibility of mental operations.

### Mental re-indexing of object positions is linked to working memory abilities

We found that pro-active swapping in context 2 was more likely when subjects also showed better performance of the delayed match-to-sample task, suggesting a relationship of working memory and pro-active swapping (**Fig. 4C**). In contrast, working memory did not correlate with the average speed of learning the sequence (**Fig. 4B**), suggesting the learning of the initial sequence does involve associative mechanisms while swapping relies on mentally manipulating a learned structure in working memory. Consistent with this suggestion the re-indexing framework postulates a transient short-term buffer of the to-be-swapped objects is needed to re-arrange the order of a sequence (Tian et al., 2024). More generally, a short-term memory buffer enables prospective planning of temporal orders of items that are not physically visible. For example, prospective working memory has been documented in NHPs using paradigms that require planning ahead by masking future items of a sequence or require comparisons of the relative rank of items that appeared in different lists (Beran et al., 2004; Inoue and Matsuzawa, 2009; Treichler and Raghanti, 2010; Scarf et al., 2011; Gazes et al., 2012; Templer et al., 2019). Our results extent these studies by suggesting that stronger working memory performance as measured with a delayed-match-to-sample-task is linked to the mental ability for flexible re-indexing of objects to non-adjacent temporal positions.

### Swapping of sequences uses long-term memory

In addition to working memory, our results also suggest that pro-active mental re-indexing is more likely when the sequential structure of object relationships has been more firmly learned in long-term memory. Subjects more likely pro-actively swapped object B and D in context 2 the better they had learned the sequence in context 1. This result suggests that pro-active swapping is a behavioral strategy that becomes available once a sequence is sufficiently well represented in memory (Jensen et al., 2021). With poorer learning of the context 1 sequence subjects applied equally likely a serial inference strategy, i.e. choosing at the second ordinal position 2 the adjacent item C instead of item D in context 2 (**Fig. 2E**). This finding extends previous studies that document monkeys represent (Chen et al., 1997; Jensen et al., 2019; Mione et al., 2020; Ferhat et al., 2022; Jensen et al., 2022), and memorize long term (Orlov et al., 2000; Templer et al., 2019) serially ordered items as their relative ordinal rank. Our results are consistent with these prior findings by suggesting that representing serially ordered objects by chaining neighboring objects and knowing their relative rank in the sequence is a ‘default strategy’ by NHPs that is applied as long as a sequence is not learned sufficiently deep, or if there is no task requirement to infer a more abstract temporal structure that would support the mental re-indexing operations of non-neighboring positions. According to this interpretation, NHPs are able to infer latent temporal structure with sufficient experience of the temporal structure and do not have a ‘hard cognitive limitation’ of inferring more complex temporal structures from item sequences beyond representing serial chains of item-item associations (Dehaene et al., 2015; Zhang et al., 2022). This conclusion is also supported by neurophysiological evidence of neurons in the prefrontal cortex as well as in the medial temporal cortex. In these brain areas neuronal responses have been found that are tuned to the ordinal rank of items in multi-item sequences (Xie et al., 2022; Chen et al., 2024a; Chen et al., 2024b; Shpektor et al., 2024). Prefrontal rank-selective neuronal responses predict the specific items that a subject uploads at each position even when that item is erroneously uploaded and lead to an unrewarded choice (Averbeck and Lee, 2007; Chen et al., 2024a). The neural coding of a rank position independently of the specific item that is encoded at that rank in principle supports a flexible assignment of items to different ranks, documenting that neuronal representations are not limited to serial item-item associations.

We speculate that the key behavioral results of our study - the ability of NHPs to flexibly re-index objects to non-adjacent positions in learned sequences - was facilitated by various design features of our task. Firstly, NHPs were exposed in each behavioral session to multiple sequence pairs with context 1 and the swapped items of context 2 already early during training. This design aspect ensured the swapping rule was not a rare exceptional task feature, but an integral part of their daily task environment, motivating them to figure out how to complete swapped sequences in context 2 in order to receive rewards. Secondly, our task paradigm enforced a ‘retouch-the-last-correct-item after-an-error’ rule, which ensured that erroneously made serial object connections were not left unnoticed by were corrected immediately by the correct pair (see **Suppl. Video**). Lastly, the task paradigm provided immediate performance feedback of every choice in the form of the halo feedback and the stepping forward or reset of the slider progress bar (**Fig. 1A**). This design aspect provided unambiguous information about erroneously chosen objects, which will have facilitated learning.

## Conclusion

Taken together, our results show that NHPs are able to flexibly re-index objects to a non-adjacent position within 5-object sequences once they have been learned at high proficiency. This mental flexibility suggests that NHPs infer latent temporal structure of their experiences when they engage in a task that requires mental operations on-top of this structure such as the required swapping of objects once the context changed. These abilities suggest that NHPs form non-spatial cognitive maps and use them to mentally manipulate items during goal-directed behavior (Whittington et al., 2022). We speculate that this mental capacity will have evolved in NHPs to support a higher level of adaptiveness of behavior during many serially organized behaviors beyond arranging visual objects in novel temporal relationships (McNamee et al., 2021).

## Acknowledgments

This work was supported by the National Institute of Mental Health (R01MH123687). The funders had no role in study design, data collection and analysis, the decision to publish, or the preparation of this manuscript.

## Data and code accessibility

Data and custom programming code for analysis is available upon request.

## Methods

### Ethic Statement

All animal and experimental procedures complied with the National Institutes of Health Guide for the Care and Use of Laboratory Animals and the Society for Neuroscience Guidelines and Policies and were approved by the Vanderbilt University Institutional Animal Care and Use Committee.

### Experimental Design

Four male rhesus macaques (Monkey S: 10 yrs / 12.6 kg; Monkey B: 10 yrs / 10.8 kg; Monkey K: 12 yrs / 11.9 kg; Monkey J: 13 yrs / 12.9 kg) were used in this study. They performed the experimental task in their housing cage using cage-mounted touch screen stations (Womelsdorf et al., 2021) (**Fig. 1A**). Visual display, behavioral response registration, and reward delivery were controlled by the Multi-Task Suite for Experiments (M-USE) (Watson et al., 2023). M-USE is an open-sourced video-engine based Unity3D platform that is integrated with a touchscreen, a video camera system and reward delivery hardware.

The task required learning the sequential temporal order of sets of five objects. We generated novel sets of objects for every new sequence by randomly assigning each object different features from up to ten different feature dimensions using multi-dimensional 3D rendered so-called Quaddle objects (Watson et al., 2019). We used Quaddle 2.0 objects that vary in ten feature dimensions (e.g. the shape, color, body pattern, different arm orientations, the presence of a head, etc.), each with >10 possible feature values (different body shapes, variable colors, etc.) (**Fig. 1C**). The objects were generated using the software Blender and custom Python scripts. For the experiment, object colors were chosen to be equidistant within the perceptually defined CIELAB color space. Objects were presented on an Elo 2094L 19.5” LCD touchscreen with a refresh rate of 60 Hz and a resolution of 1,920 × 1,080 pixels, rendered at 2.9-4.2 × 2.4-4.7 cm on the screen.

For each sequence, a new contextual background image was displayed. Context images were generated with DALL·E 2, an AI system developed by OpenAI, using a text prompt for obtaining images from the categories rocks and sand. We applied color filters to images to obtain a larger number of distinct context backgrounds, ensuring each color filter maintained a fixed luminance value of 50. The colors were selected by evenly spacing them around the CIELAB color wheel, ensuring a diverse range of hues. The brightness of the background images was adjusted to be ≤ 50% of the HSL scale.

### Task Paradigm

On each trial six objects were presented at a random location at equal eccentricity relative to the center of the screen (**Fig. 1A**). Five of the objects were assigned a unique ordinal temporal position in the sequence, while a sixth object was a sequence-irrelevant distractor. Each sequence was presented for a maximum of 15 trials. In each trial subjects had a up to 15 choices to complete the sequence and earn fluid reward by touching objects and receiving either positive feedback (a yellow halo and high pitch sound) for correct choices, or negative feedback (a transient grey halo and low pitch sound) for touching an object at an incorrect temporal position. After an erroneous choice, subjects had to re-choose the last correct object in the sequence before searching for the next object in the sequence (see **Suppl. Video**). When a trial was completed the objects were removed from the screen and a new trial was started with the objects displayed at random new locations equidistant from the center of the display. For each correctly chosen object, the slider position of the slider progress bar on top of the screen stepped forward. Successful completion of a sequence always completed the slider progress bar and resulted in a water reward. When a sequence was not completed no fluid reward was given and the slider progress bar was reset for the subsequent trial with the same objects at new random locations. After completing fifteen trials on a sequence A-B-C-D-E we changed the background context and swapped the order of object B and D, requiring to perform the sequence A-D-C-B-E for the next fifteen trials (**Fig. 1C**). We refer to the initial and the swapped sequence as a ‘sequence pair’.

### Delayed presentation of novel and familiar sequences and same or different contexts

Each session began by presenting four or five sequence pairs in a first set, followed by interleaving 120 trials of a delayed match to sample task, and followed by a second set with five or six sequence pairs (**Fig. 1B**). In this second set one of the sequence pairs was novel, and four or five pairs were repeated sequence pairs from the early first set of sequences presented prior to the delayed match-to-sample task (**Fig. 1B**). Fifty to seventy-five percent of sequences in the repeated set were presented with the same context, while the remaining sequences were randomly presented with different contexts.

### Distractor objects

In each trial, one of six objects was a sequence-irrelevant distractor. Feature dimensions of the distractor were chosen to have a high degree of feature similarity to the second object of the initial sequence (which then became the fourth object in the swapped sequence). The sequence-irrelevant distractor object differed from the sequence relevant object at ordinal position two (object B) in only three features, while the other objects of the sequence did not share features (**Fig. 1C**). The distractor object was used to validate subjects paid attention to the features of the objects. Analysis results (**Suppl. Fig. S5**) confirmed that subjects indeed confused the distractor object most likely with object B in the first sequence in context 1 (2^nd^ ordinal position) and also with object B in the second swapped sequence in context 2 (4^th^ ordinal position).

### Interleaved delayed-match-to-sample working memory task

After the first and before the second set of sequences, subjects performed a delayed match-to-sample task (DMTS) for 120 trials (**Fig. 4A**). The DMTS task is part of the M-USE platform (Watson et al., 2023). DMTS used Quaddle 1.0 stimuli that varied in up-to four feature dimensions (body shapes, arm style, body pattern, color). The DMTS trial presented a sample object for 0.5 s, followed by a delay of 0.5, 1.25 or 1.75 seconds, before two or three test objects were shown. One of the test objects matched the sample and when touched resulted in a yellow halo, a high pitch sound and a token reward (a green circle) that was added to a token bar (Watson et al., 2023). Choosing non-matching objects resulted in a grey halo, a low-pitched tone, and a grey token that was subtracted from tokens available in the token bar. The token bar contained three placeholders for tokens and flashed white/red when all three tokens were completed, resulting in the delivery of fluid reward.

### Analysis of sequence learning

In each trial subjects could make maximally ten errors to reach the last, fifth object in a sequence or else a new trial started with the same objects in new positions. Up to fifteen trials were shown with the same sequence. We quantified learning as the proportion of trials in which an object sequence was completed (**Fig. 1C**). A sigmoid fit was applied to the completion rates across trials from all subjects. The trial at which learning reached criterion performance was defined as the first trial at which subjects completed 80% of trials correctly (**Fig. 1C**). To assess overall performance, we calculated the average completion rate for each subject’s sessions. We defined well performed sequences when more than 80% of the trials were completed; blocks falling below this threshold were considered poorly performed (**Suppl. Fig. 1D**). To evaluate the consistency of learning at each ordinal position we calculated the proportion of correct choices for objects at each ordinal position across trials within a block (**Fig. 1D**). For each ordinal position, we identified the trial at which subjects achieved 80% correct object choices (**Fig. 1E**).

To analyze how learning over trials correlated with changes in choice reaction times, we calculated the average time of subsequent choices for each ordinal position (**Suppl. Fig. 2**). We plotted the reaction times separately for trials before and after reaching the trials-to-80% criterion completion rate to investigate the effect of learning on reaction time. A Welch t-test was applied to compare the reaction time differences before and after the learning point at each ordinal position.

### Analysis of errors

To evaluate how subjects learned, we quantified the decrease in the proportion of different error types across trials (**Suppl. Fig. 1D-F**): (1) An exploration error occurred when choosing an incorrect object among objects not yet learned; (2) A rule-breaking error involved incorrectly re-selecting or failing to re-select the last correctly chosen object after an error, or re-selecting a previously correctly chosen object; (3) A distractor error occurred when choosing the distractor object; (4) A perseverative error involved choosing the same incorrect object as in the last choice before reselection. We calculated the overall proportion of each error type relative to all errors as a function of the trial in a block. To evaluate how fast subjects reduced each type of error over trials we fitted an exponential decay function (*a* ∗ exp(−*b* ∗ *x*) + c) across fifteen trials. The same decay function was applied to each session individually, and decay factors were plotted to compare the decreasing speed of error types (**Suppl. Fig. 1D**). Statistical analysis was performed using one-sample t-tests to compare the decay factor to zero, and a Welch t-test was applied to compare each pair of error types.

### Analysis of re-ordered sequence with swapped object positions

We tested how subjects learned the swapping of objects B and D from the first sequence A-B-C-D-E in context 1 to the second sequence A-D-C-B-E in context 2. A sigmoid fit was applied to the completion rates over trials in context 1 and context 2 (**Fig. 2B**). We calculated the average trials needed to reach learning criterion (≥80% completion rate) for each session across sequence pairs in context 1 and context 2, using a t-test to compare the differences (**Fig. 2C**). To investigate the effect of swapping on search time, we plotted the average reaction time for each ordinal position in both conditions (**Suppl. Fig. S2B**). Next, we analyzed which objects subjects chose after they incorrectly chose object B at the 2^nd^ ordinal position in context 2 (**Fig. 2D**). A t-test was used to compare the transition probability of objects C, D, or E. We also calculated the probabilities of choice separately for well-performed and poorly performed blocks (**Fig. 2E**). To quantify how likely subjects chose objects at each ordinal position in context 1 and context 2 before they reached the 80% learning criterion, we calculated transition choice probabilities after choosing object A (**Fig. 2F**) and after choosing correctly object D in context 2 (**Fig. 2G**). We used t-tests to compare whether subjects inferred the correct swapped order in context 2 after choosing object D at the 2^nd^ ordinal position (choosing C), or whether they confused the sequential ordering in context 2 with the order from context 1 (and incorrectly chose E).

### Analysis of effect of memory on swapping

*To quantify the effect of memory on sequence learning* we tested whether sequences that were repeated late in the session were performed better than novel sequences performed late in the session. Using paired t-tests, we compared how fast sequences were performed at learning criterion (≥80% completion) that were shown early (‘new early’), or repeated late in the session (‘repeated’). We also included novel sequence pairs that were presented late in the session (‘new late’), or introduced newly late in the session (‘new late’) (**Fig. 1B**; **Fig. 3A**). Next, we analyzed how memory influenced pro-active swapping and retro-active swapping behavior. We calculated two measures: a value indexing the likelihood that subjects chose object C instead of object D in context 2 at the second ordinal position (an incorrect serial inference), defined as: *p*(choosing *D* − choosing *C* | erroreously choose *B*; and we calculated a value indexing the likelihood of choosing object D at the second ordinal position in context 2, reflecting pro-active swapping: *p*(choosing *D* − choosing *B* | correctly choose A). T-tests were used to quantify the difference of these strategies in these repeated versus new/late new sequence pairs (**Fig. 3B,C**).

### Analysis of the effect of learning proficiency and task experience on swapping behavior

We quantified how the proficiency of performing a sequence (the average completion rate in context 1) affected the likelihood of pro-active swapping behavior in context 2. We used t-tests to compare the values indexing the likelihood of showing incorrectly a serial inference strategy or a pro-active swapping strategy in context 2 for sequences performed at low versus high completion rates in context 1 (**Fig. 3D,F**). We also correlated these values on a sequence-by-sequence basis using linear regression (**Fig. 3E,G**) and used linear regression of the performance across sessions to discern the influence of experience over the course of the experiment (**Fig. 3H,I**).

### Analysis of the relation of working memory and sequence learning

The delayed match-to-sample task presented a sample stimulus, followed by a blank delay lasting 0.5, 1.25, or 1.75s, followed by the presentation of three objects (2 distractors and 1 target object matching the sample stimulus) (**Fig. 4A**). The distractor stimuli were either similar or dissimilar to the target stimulus, because they shared, or did not share, features with the target objects. We analyzed the overall accuracy for low/high similarity and across delays. Linear regression analysis showed no decline across delays so that we averaged the accuracy across conditions for individual sessions and each monkey (**Suppl. Fig. 4**). To analyze the relationship of working memory, sequence learning and the ability to anticipate the swapped object order in context 2, we applied individual linear regressions and a linear mixed model fit to the performance across sessions (**Fig. 4B,C**).

### Analysis of context repetition on sequence learning

In a subset of sequence pairs shown late in the session, we repeated the object sequences from early in the session but on a different background context. To test how the difference of contexts versus the similarity of contexts affected performance and the likelihood to observe pro-active swapping behavior in context 2 we determined the difference in accuracy between early and late (repeated) sequences with the same and different contexts (**Fig. 4D-F**). Paired t-tests were used to compare the accuracy between conditions.

### Analysis of choices of the distractor

For very 5-object sequence the task presented a sixth distracting object that was irrelevant to the sequences and shared three features with the object B. We tested whether the distractor object was more likely chosen at the ordinal position at which it shared features with object B of the sequence than at other ordinal positions. We tested for differences in the proportion of distractor choices in context 1 and 2 using a proportion Z-test (**Suppl. Fig. S5**).

