## Supplementary Figures for "Flexible Learning and Re-ordering of Context-dependent Object Sequences in Nonhuman Primates"

### Supplementary Figures S1-S5

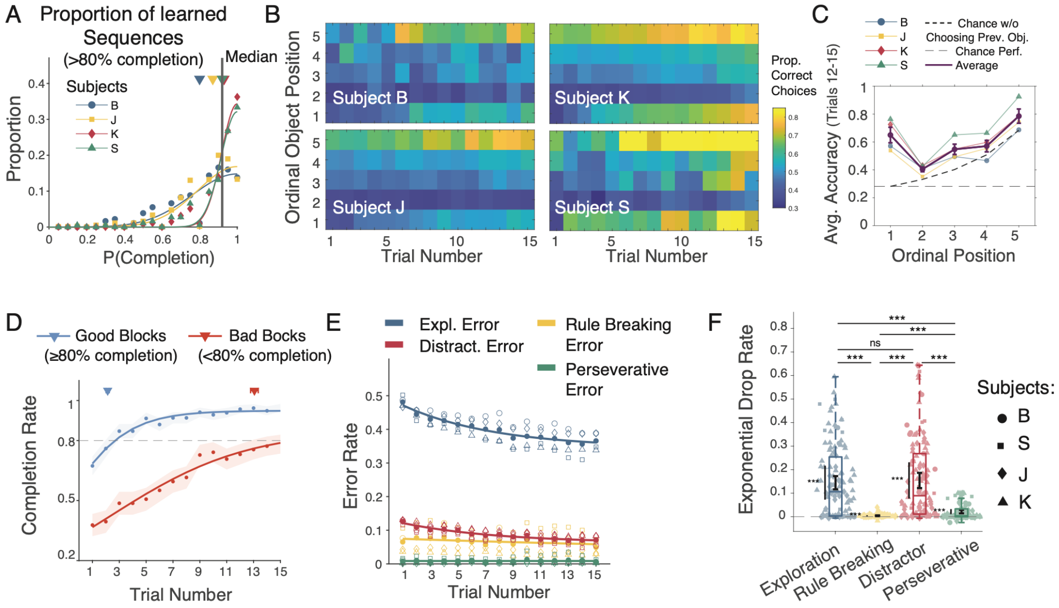


**Supplementary Figure S1. Overall learning performance.** (**A**) Prop. of sequences at which subjects reached >80% completion. (**B**) Prop. correct choices for each ordinal position (y-axis) and trial (x-axis). (**C**) Prop. of correct choices for each ordinal position averaged across last 4 trials. Error bars are SE's. Means (± SE) were: ordinal position 1: 0.6501 ± 0.0556, position 2: 0.4049 ± 0.0180; position 3: 0.5475 ± 0.0369; position 4: 0.5706 ± 0.0413; position 5: 0.7858 ± 0.0509. The darker dashed line indicates chance accuracy expected without reselecting previously correct objects, while the lighter dashed line represents chance accuracy when reselecting previously correct objects is allowed. (**D**) Avg. completion rates for blocks with >80 and ≤80% completion rate. (**E**) Exploration errors, distractor errors, and rule-breaking errors decreased over trials. Data were fit with an exponential decay function ($y = a * \exp(-bx) + c$). (**F**) Mean (and 95% CI) of the decay factor (b) for each error type: Exploration error (0.1447 [0.1167, 0.1727]), Rule breaking error (0.0041 [0.0025, 0.0058]), Distractor error (0.1543 [0.1220, 0.1867]), and Perseverative error (0.0216 [0.0145, 0.0288]). Exploration and distractor errors had a similar decay rate (p = 0.656). Stars denote significance levels for pairwise comparisons (Welch's t-tests). Decay factors for all error types were significantly greater than zero, confirming a decrease in all error types over the course of learning (t-test against zero, p-values: Exploration error: 6.6 × 10^-18; Rule breaking error: 5.4 × 10^-6; Distractor error: 7.3 × 10^-16; Perseverative error: 4.3 × 10^-8).

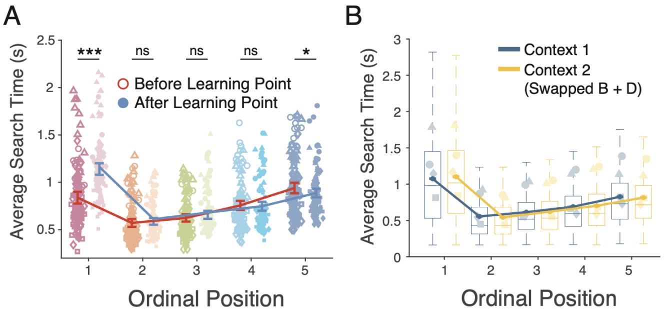


**Supplementary Figure S2. Reaction times to objects across ordinal positions during and after sequence learning.** (**A**) Average reaction times for correct touches before and after learning point. Before learning points, ordinal position 1 to 5 respectively (Mean ± 95%CI): 0.84 ± 0.06, 1.14 ± 0.06, 0.57 ± 0.04, 0.59 ± 0.04, 0.62 ± 0.04; After learning points: 0.66 ± 0.04, 0.76 ± 0.05, 0.72 ± 0.05, 0.94 ± 0.06, 0.86 ± 0.05. Welch's t-test was applied to each ordinal position, and significant difference was found at the first and the last ordinal positions (Position 1: $p < 0.00001$, Position 2: $p = 0.5520$, Position 3: $p = 0.2550$, Position 4: $p = 0.3416$, Position 5: $p = 0.0384$). (**B**) Average reaction times for correct touches before (blue) and after swap (yellow) for each subject. No significant difference was found in all ordinal positions.

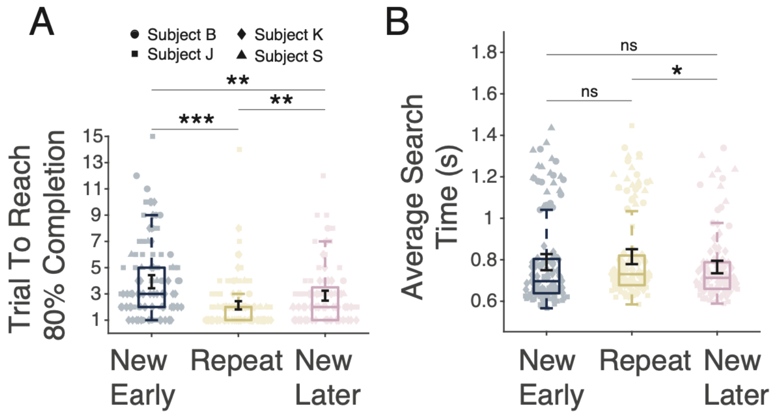


**Supplementary Figure S3. Memory of sequences improved performance.** (**A**) Sequences repeated after the working memory task are performed better. In the new early condition, the 80% completion rate was reached after 3.93 ± 0.50 trials (Mean ± 95%CI), whereas, in the repeat condition, it was significantly quicker at just 2.13 ± 0.32 trials (New early vs Repeat: p: 1.19 × 10^-8). For new sequences introduced late (to control for 'time in the session'), the avg. trial to reach the 80% criterion was 2.87 ± 0.38, demonstrating that while there was some improvement in performance for new sequences in the later set, the most substantial gain was observed for the repeated sequences (New early vs New late: p: 0.0014; Repeat vs New late: p: 0.0052). (**B**) Choice reaction times for correct choices at each ordinal position were calculated to check if there were any difference in three conditions (New early: 0.79 ± 0.04; Repeat: 0.81 ± 0.04; New late: 0.77 ± 0.031). We found a slightly slower reaction time for the repeat blocks when compared with New late blocks (p: 0.039), but no other comparisons were significant (New early vs Repeat: p: 0.33; New early vs New late: p: 0.35).

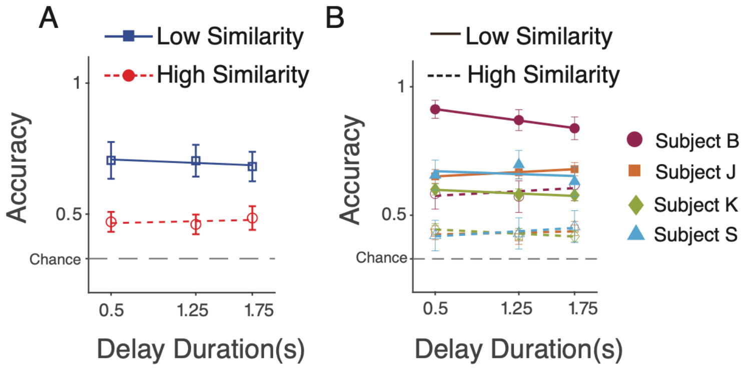


**Supplementary Figure S4. Working memory (delayed match-to-sample) performance.** (**A**) Task performance across varying object feature similarities and delay times. Chance performance was set at 0.33. Accuracy was significantly higher for the Low Similarity condition compared to the High Similarity condition. High Similarity condition: Intercept: 0.4606; Slope: 0.0097; p = 0.6670; Low Similarity condition: Intercept: 0.7172; Slope: -0.0175; p = 0.3772. The overall average accuracy across all conditions was 0.59 ± 0.16 (Mean ± 95% CI), notably above chance (0.33). For the 0.50s delay condition, performance was 0.47 ± 0.12 (High Similarity) and 0.71 ± 0.22 (Low Similarity); for the 1.25s delay condition, it was 0.46 ± 0.12 (High Similarity) and 0.70 ± 0.19 (Low Similarity); for the 1.75s delay condition, 0.48 ± 0.14 (High Similarity) and 0.68 ± 0.18 (Low Similarity). (**B**) Individual subject performance across all similarity and delay time conditions. Subject-level data remained consistent, with similar patterns observed between high and low similarity conditions, regardless of delay time.

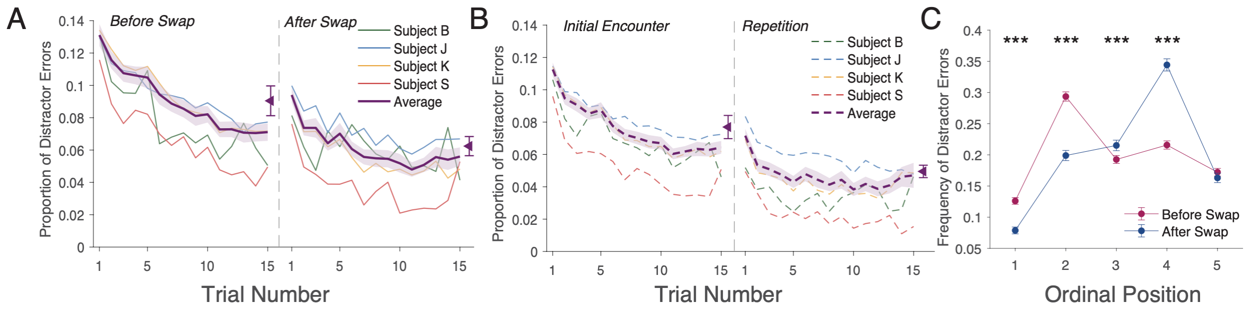


**Supplementary Figure S5. Learning to ignore distractor object. (A)** Proportion of distractor choices across trials in context 1 (*left* panel) and context 2 (*right* panel). The mean value is shown as the rightmost data point (Mean ± 95% CI, Before swap: 0.090 ± 0.010; After swap: 0.061 ± 0.006). **(B)** Same format as A for distractor choices in sequences early (left) and late (right) in the session. The average is shown as rightmost data point (Mean ± 95% CI, Initial encounter: 0.076 ± 0.008; Repetition: 0.046 ± 0.004). **(C)** Distractor choices at each ordinal position in context 1 (distractor is similar to B at the 2$^{nd}$ ordinal position) and in context 2 (distractor is similar to B at the 4$^{th}$ ordinal position). Two-proportion Z-tests were applied at each ordinal position for comparing the difference. Stars denote sign. level. (Ordinal Position: Before & After Swap, Mean ± 95%CI) 1: 0.13 ± 0.01; 1: 0.08 ± 0.01;  2: 0.29 ± 0.01; 2: 0.20 ± 0.01; 3: 0.19 ± 0.01; 3: 0.22 ± 0.01; 4: 0.22 ± 0.01; 4: 0.34 ± 0.01; 5: 0.17 ± 0.01; 5: 0.16 ± 0.01.
